# TMExplorer: A Tumour Microenvironment Single-cell RNAseq Database and Search Tool

**DOI:** 10.1101/2020.10.31.362988

**Authors:** Erik Christensen, Alaine Naidas, Mia Husic, Parisa Shooshtari

## Abstract

Tumour microenvironments (TME) contain a variety of cells including but not limited to stromal fibroblasts, endothelial cells, immune cells, malignant cells, and cells of the tissues of origin, whose interactions likely influence tumour behaviour and response to cancer treatment. The specific composition of the TME can be elucidated using single-cell RNA sequencing (scRNA-seq) by measuring expression profiles of individual cells. Several scRNA-seq datasets from multiple cancer types have been published in recent years, yet we still lack a comprehensive database for the collection and presentation of TME data from these studies in an easily accessible format. We have thus built a database of TME scRNA-seq data, containing 21 TME scRNA-seq datasets from 12 different cancer types. We have also created an R package called TMExplorer, which provides an interface to easily search and access all available datasets and their metadata. Data and metadata are kept in a consistent format across all datasets, with multiple expression formats available depending on the use case. Users can view a table of metadata and select individual datasets or filter them by specific characteristics. Users may also select a specific type of cancer and view all published scRNA-seq data for that cancer type available in our database. Users are provided with an option to save the data in multiple formats in order to view or process it outside of R. Thus, the TMExplorer database and search tool allows for thorough examination of the TME using scRNA-seq in a way that is streamlined and allows for easy integration into already existing scRNA-seq analysis pipelines.

## INTRODUCTION

Single-cell RNA sequencing (scRNA-seq) is a new technology that has emerged as an important tool to measure gene expression for individual cells, enabling the examination of cellular heterogeneity and tissue composition with incredible precision. This has been particularly applicable in cancer research for the study of tumour composition, heterogeneity and phenotype, all of which are directly impacted by the tumour microenivronment (TME). TMEs are composed of different stromal and cancer cell types whose interactions likely dictate different aspects of tumour behaviour, such as metastasis (1–4). Combined with scRNA-seq analysis methods, scRNA-seq enables us to dissect the TME into individual cells and investigate the different cell subpopulations that exist. Such investigations into the TME are becoming increasingly important, as tumour composition and heterogeneity can influence cancer progression and the outcome of cancer therapy (1, 4–9).

With the advancement of scRNA-seq in cancer research, the number of TME datasets that are generated continues to increase, yet they can be difficult to access. Raw sequence reads generated by scRNAseq can be shared through online archives, such as the Sequence Read Archive (SRA) (10), however they exist as large files that require further processing to be analyzed, making data access a challenge. Already processed scRNA-seq data containing gene expression information can be accessed through online archives, such as the Genome Expression Omnibus (GEO) (11), and can be more easily downloaded for use in one’s own analysis. Furthermore, to manage the growing abundance of publicly available scRNA-seq data, proper quality control and curation of datasets must be done (12, 13). Currently, several online databases offer curated collections of public scRNA-seq datasets, such as PanglaoDB (12), scRNASeqDB (13), JingleBells (14) and the Single Cell Portal created by the Broad Institute of MIT and Harvard (https://singlecell.broadinstitute.org/single_cell) (15). Most existing scRNA-seq databases include a mixture of samples from normal tissues and tissues affected by cancer or other diseases (12–14), while others focus primarily on samples from normal tissues (16, 17). A recently published toolkit called CReSCENT (18) contains only cancer scRNA-seq data, however it mainly acts as a cancer data analysis pipeline rather than a database. A comprehensive database for the collection and sharing of TME scRNA-seq datasets from a range of tumour types does not yet exist, and researchers interested in using publicly available TME data must search through several databases to collect relevant datasets for their study. A database of TME scRNA-seq samples will thus streamline the data collection steps required for researching cancer at a single-cell level, lowering the barrier for entry to this type of study.

It is likewise important that scRNA-seq databases are designed to facilitate streamlined data collection and analysis. This can include a search tool that allows users to select datasets based on desired characteristics. While existing databases include search tools, they provide few options in characteristics users are able to search for and often require users to browse through a metadata table prior to selecting datasets of interest. Furthermore, they are designed as webbased tools, and thus are not intended to be integrated into workflows (12–15). Workflow integration would enable users to access data directly in their pipelines, thus automating the data collection process and increasing analysis efficiency. A scRNA-seq database that is provided as an R-package and contains a comprehensive search tool which allows users to select datasets based on a wider variety of characteristics would make the data collection process easier for researchers.

Here, we present a curated collection of tumour scRNA-seq datasets made available as an R-package called TMExplorer. TMExplorer contains publicly available scRNA-seq datasets specific to TMEs from various tumour types collected from different scRNA-seq studies (1–3, 5–9, 19–31) and online databases (11, 32). In addition to gene expression data, TMExplorer contains the corresponding cell type annotations and gene-signature information for several datasets, and provides a search tool enables users to search for multiple datasets according to 13 different characteristics (Table 1). When selecting datasets, users can review the metadata table first or they can retrieve datasets that match specific criteria without having to browse through the metadata table. While online databases require users to download a given dataset prior to use, TMExplorer allows users to access and search available datasets within R. Users can thus input the data directly into existing pipelines with only a few commands. Each dataset can be used directly within R as a *SingleCellExperiment* object, or exported as a gene expression matrix in multiple formats for use with other applications. Users interested in validating scRNA-seq analysis algorithms, as they apply to TME data, can easily access this information through TMExplorer and incorporate it into their pipelines. Altogether, TMExplorer makes it easier for researchers to access and share TME scRNA-seq datasets, facilitating the study of TMEs at the single-cell level in the field of cancer research.

**Table 1:** A list of search parameters that can be passed to queryTME in order to filter the available datasets.

| Search Parameter | Description |
| --- | --- |
| geo_accession | Search by GEO accession |
| score_type | Search by type of score available |
| has_signatures | Search by presence of cell type signature gene sets |
| has_truth | Search by presence of cell type annotations |
| tumour_type | Search by type of tumour |
| author | Search by first author |
| journal | Search by publication journal |
| year | Search by publication year |
| pmid | Search by PMID |
| sequence_tech | Search by sequencing technology |
| organism | Search by source organism |
| sparse | Return expression in sparse matrices |
| download_format | Specify a list of score formats to download.<br>Additional formats will be stored in altExps |

## METHODS

### Data Collection

In order to collect the datasets, we searched the National Center for Biotechnology Information (NCBI) (33) for relevant scRNA-seq studies using the following keywords: single cell RNA sequencing, tumour, cancer, tumour microenvironment, and malignant. We then carefully reviewed the published literature and any associated data to confirm if they matched our criteria. Datasets were included in our data collection if they were publicly available as processed data, were generated by scRNA-seq and if they consisted of TME expression data. A total of 21 datasets originating from different types of human and mouse tumours were collected from online sources such as the NCBI’s Gene Expression Omnibus (GEO) (11), ArrayExpress (32), and Github (34). Out of the 21 datasets we collected, 18 datasets originated from human tumours and 3 datasets originated from mouse tumours. Descriptions of the collected datasets are provided in Table 2. Metadata for each dataset, such as tumour type and number of cells sequenced, were collected from descriptions in the corresponding publications and/or from the online sources that the datasets were obtained from. If publicly available, we also retrieved cell-type annotations and/or gene signature information that accompanied the datasets.

**Table 2:** List of tumor microenvironment scRNA-seq datasets included in TMExplorer

| **Dataset** | **Cancer type** | **Sequencing Technology** | **Number of tumors** | **Number of cells** | **Number of genes** | **Annotation available?** | **Gene signature available?** |
| --- | --- | --- | --- | --- | --- | --- | --- |
| **Patel *et al.* Science 2014** | Primary Glioblastoma | SMART-seq | 5 human primary glioblastoma tumors | 1,456 | 5,796 | Yes | No |
| **Tirosh *et al.* Science 2016** | Metastatic melanoma | SMART-seq2 | 19 human melanoma tumors | 4,645 | 23,686 | Yes | Yes |
| **Tirosh *et al.* Nature 2016** | Oligodendroglioma | SMART-seq2 | 6 human oligodendroglioma tumors | 4,347 | 23,686 | Yes | Yes |
| **Venteicher *et al.* Science 2017** | IDH-mutant astrocytoma | SMART-seq2 | 10 human IDH-mutant Astrocytoma tumors | 6,341 | 23,686 | No | No |
| **Li *et al.* Nature Genetics 2017** | Colorectal cancer | Fluidigm C1 | 11 human primary colorectal cancer tumors | 376 | 57,241 | Yes | Yes |
| **Chung *et al.* Nature Commun 2017** | Primary breast cancer | Fluidigm C1 | 11 human primary breast cancer tumors | 564 | 57,915 | Yes | Yes |
| **Puram *et al.* Cell 2017** | Head and neck squamous cell carcinoma | SMART-seq2 | 18 human primary oral cavity tumors and 5 lymph node metastases | 5,902 | 21,884 | Yes | Yes |
| **Giustacchini *et al.* Nature Med 2017** | Chronic myeloid leukemia | SMART-seq2 | 20 human bone marrow aspirates | 2,287 | 23,384 | No | No |

| Dataset | Cancer type | Sequencing Technology | Number of tumors | Number of cells | Number of genes | Annotation available? | Gene signature available? |
| --- | --- | --- | --- | --- | --- | --- | --- |
| **Filbin *et al.* Science 2018** | H3K27-mutant glioma | SMART-seq2 | 6 human primary H3K27M-glioma tumors | 4,059 | 23,686 | Yes | No |
| **Jerby-Arnon *et al.* Cell 2018** | Melanoma | SMART-seq2 | 33 human melanoma tumors | 7,186 | 23,686 | Yes | Yes |
| **VanGalen *et al.* Cell 2019** | Acute myeloid leukemia | Seq-well | 40 human bone marrow aspirates | 38,410 | 27,899 | No | No |
| **Ting *et al.* Cell Reports 2014** | Pancreatic circulating tumor cells | Tang protocol | 5 mice with pancreatic cancer, 1 mouse embryonic fibroblast cell line, 1 mouse pancreatic cancer cell line, 1 control mouse | 187 | 29,018 | No | No |
| **Miyamoto *et al.* Science 2015** | Prostate circulating tumor cells | ABI SOLiD | 18 patients with metastatic prostate cancer, 4 patients with localized prostate cancer, 12 bulk primary prostate tumors, 4 prostate cancer cell lines | 169 | 21,696 | No | No |
| **Jordan *et al.* Nature 2016** | Breast circulating tumor cells | SMARTer | 2 ER+/HER2- breast cancer patients, 14 triple negative breast cancer patients | 74 | 23,368 | No | No |
| **Azizi *et al.* Cell 2018** | Breast carcinoma | InDrop | 8 human breast carcinomas | 46,016 | 14,875 | No | No |
| **Lambrechts *et al.* Nature Med 2018** | Non-small cell lung carcinoma | 10x Genomics | 5 human non-metastatic lung squamous carcinoma tumors | 51,775 | 22,180 | Yes | Yes |
| **Davidson *et al.* bioRxiv 2018** | Melanoma | SMART-seq2 | ? mouse tumors | 6,423 | 26,946 | No | No |
| **Peng *et al.* Cell Research 2019** | Pancreatic ductal adenocarcinoma | 10x Genomics | 24 human primary pancreatic ductal adenocarcinoma tumors, 11 control pancreases | 57,530 | 24,005 | Yes | Yes |
| **Darmanis *et al.* Cell Reports 2017** | Primary glioblastoma | SMART-seq2 | 4 human glioblastoma tumors | 3,589 | 23,465 | No | No |
| **Kumar *et al.* Cell Reports 2018** | Mixed cancer: Melanoma, breast mammary carcinoma, Lewis lung carcinoma, colon carcinoma, fibrosarcoma | 10x Genomics | 1 mouse melanoma tumour, 1 mouse breast mammary carcinoma tumour, 1 mouse Lewis lung carcinoma tumor, 2 different mouse colon carcinoma tumors, 1 mouse fibrosarcoma tumor | 10,573 | 27,998 | No | No |
| **Zhao *et al.* BMC Med Genomics 2019** | Glioblastoma cell line | Fluidigm C1 | 1 human glioblastoma cancer cell line, 1 normal neural stem cell line | 134 | 21,209 | No | No |

### Data Curation

Datasets found on GEO often contain extra information such as Ensemble ID or chromosome region in additional rows or columns. We modified all datasets using R to ensure they followed a similar genes-by-cells format with the gene column serving as an index. Having a similar format for datasets reduces the preprocessing required to use this data in other analysis pipelines.

There are three main components to each dataset in our database: gene-by-cell expression matrices, cell type labels, and gene signatures. The cell type labels are R *dataframes* with two columns; one contains every cell barcode present in the expression matrix, the other one contains that cell’s type. The gene signatures are stored in R *dataframes* containing one column per cell type, with a list of genes that are differentially expressed by that cell type and reported in the original paper in which each dataset was first introduced. All data for each dataset is accessible within a single object in order to make it as easy to use as possible.

Since R BioConductor has existing infrastructure for working with scRNA-seq data (35, 36), we used it as the platform to build our package upon. In order to maintain compatibility with existing Bioconductor software, we return all datasets as *SingleCellExperiment* objects (36). Figure 1 shows the structure of a *SingleCellExperiment* object, where the expression data is stored as a named assay, cell type labels (if present) are stored under *colData()*, and all other information is stored in a metadata list.

- **Expression Data:** Named assays allow certain formats to be easily accessed with *getter* functions such as *counts()* and *tpm()*, while other formats can still be accessed with the *assay() getter* function (36). All *SingleCellExperiments* have one assay named according to the type of score (e.g. Counts and TPM) represented in that object. Calling *assay()* returns an expression matrix with rows of genes and columns of cells.
- **Cell Type Labels:** *ColData* stores metadata for the columns in the assay matrix. In our case this refers to the cell type annotations, if they are available. *ColData* is a dataframe that always has one row for every column in the assay matrix, ensuring that there is a label present for every cell. If the cell type is not available for a given cell, it is labelled as “unknown”.
- **Metadata:** The metadata list serves to store any other information that does not fit into a pre-existing attribute of the *SingleCellExperiment* object, and is accessed with the *metadata()* function. This named list contains the signature gene sets, available score types, tumour and host organism type, sequencing technology, author, and all other descriptive information as strings. All information that is available in the metadata table can be accessed by calling the query function of TMExplorer (i.e. *queryTME*) with the *metadata_only* parameter set to true.

**Figure 1:**
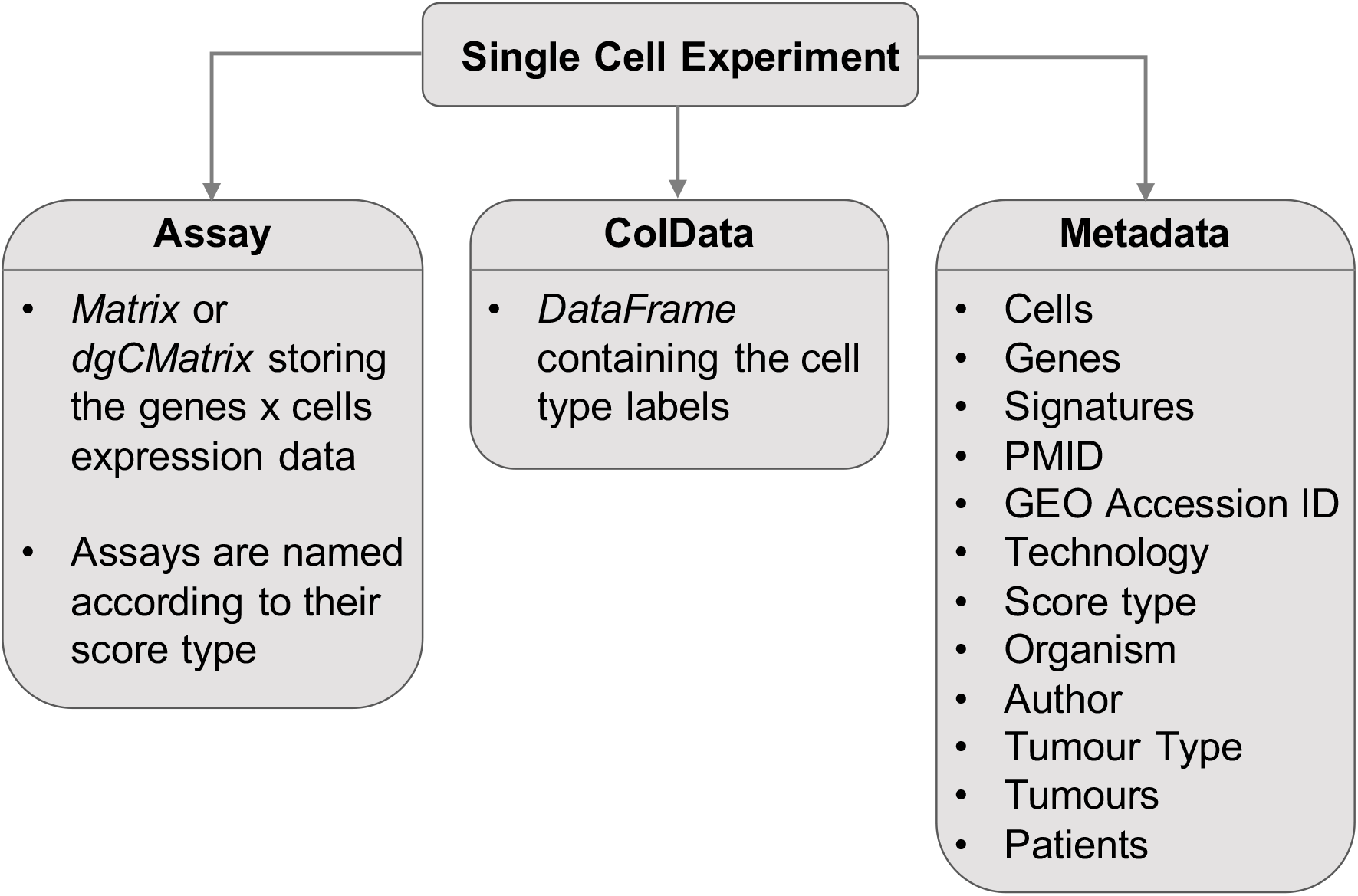
The format of the *SingleCellExperiment* objects containing TME datasets. The Assay is a matrix or *dgCMatrix* containing the gene expression table, named according to the type of score (i.e. an Assay containing raw counts would be named “Counts”); *colData* is a *DataFrame* with the number of rows equal to the number of columns in the Assay and describes the cells in the dataset; Metadata is a named list of additional metadata objects describing the dataset. A *SingleCellExperiment* object may contain one or more *AltExps*, which are nested *SingleCellExperiment* objects containing a different score type in the Assay.

### Metadata

After collecting the datasets, corresponding metadata was compiled into a table which serves as the core of the package. The metadata table contains information such as GEO accession, author, journal, year, PMID, sequencing technology, expression score type(s), source organism, type of cancer, number of patients, tumours, cells and genes, and the database that the data was obtained from. All items in the metadata table were chosen as either entities that distinguish one dataset from the others or criteria that may make a dataset or group of datasets interesting to researchers (e.g. a specific tumour type or availability of cell labels or gene signatures). Users can view the available data using the metadata table and decide which dataset best fits their needs.

### Database Query

TMExplorer provides a query function (i.e. *queryTME*) that users can employ in order to select multiple datasets based on their desired characteristics. For example, users can select specific studies by PMID or GEO accession, or filter subsets by sequencing technology, whether cell type labels or cell type signature gene sets are available etc. Sequencing technology, score type, organism, tumour type, and year were all chosen as search parameters because they represent differences in the type of data and make it easier to find data that fits the needs of different studies. We have also made it possible to search for datasets for which the cell labels and gene signatures are available. This facilitates developing and testing algorithms that require specific types of dataset information. For example, testing cell classification algorithms requires cell labels that can be used as a gold standard, and many existing algorithms require gene signatures that represent the cell types in the dataset (37, 38).

### Alternative Experiments

For several datasets, gene expressions are available in multiple score types including raw counts and normalized data by FPKM, TPM or CPM. In order to store each dataset in multiple score types, we used nested *SingleCellExperiments* objects with the alternative experiments (*altExps*) concept. Alternative experiments are guaranteed to have the same dimensions as the primary object, but can be kept separate for use in other pipelines (36). This allows users to download multiple types of scoring for use in different steps of analysis while still being able to access each dataset through a single object. Being able to download multiple score types allows our datasets to be used in a variety of algorithms that require a specific type of score, and keeping them separated as nested objects prevents accidentally applying an algorithm to the wrong score type.

### Dense vs. Sparse Data Formats

In order to reduce the memory requirements for working with large datasets, expression data is optionally available as a sparse matrix. We implemented sparse matrices using the *dgCMatrix* class from R Matrix (39). This reduces memory usage by only storing non-zero expression values. With sparse matrices, the memory required to store a dataset is reduced by as much as 8 Gb for a dataset with 51,775 cells and 22,533 genes. It should be noted that not all software packages are compatible with sparse matrices, and converting large datasets from sparse to dense may crash R on machines with low memory. Thus users should confirm that their algorithms support sparse matrices before using them. By default, TMExplorer returns dense matrices to avoid these problems.

### Exporting Data in Multiple Formats

Several tools for scRNA-seq analysis are written in R and therefore a *SingleCellExperiment* object can easily be incorporated into these pipelines and tools. However, many other analysis tools are written in Python or as webapps (18, 40, 41). To facilitate the use of TMExplorer with these tools, we wrote a function *saveTME* that writes individual TME datasets to disk as CSV or Matrix Market files, depending on whether data was loaded as dense or sparse matrices by *queryTME*, respectively. *SaveTME* takes a *SingleCellExperiment* object and a path to an output directory as parameters and saves the gene expression matrix, cell type labels, and cell type signature gene sets to disk. The resulting files can then be converted as needed and used in other applications.

## RESULTS

### Overview of the TMExplorer Package

To make it as easy as possible to integrate TMExplorer into other pipelines, all interactions with the package are done directly in R. Here, the *queryTME* function serves as the primary interface for the package, allowing users to view the metadata for all available datasets, or select a subset of datasets according to descriptive criteria (Figure 2a). *queryTME* provides a set of parameters (Table 1) used to select a subset of datasets according to characteristics. To review the available datasets, the *metadata_only* parameter should be set to *TRUE* when querying the package, and a table describing the datasets will be returned instead of the datasets themselves. The search parameters can be used to find relevant data without requiring users to review the metadata table first, lowering the barrier for use. For example, users looking for a certain type of cancer, such as melanoma, can search using *queryTME*(*tumour_type=“Melanoma”*) without needing to first examine the metadata for datasets contain melanoma cancers.

**Figure 2:**
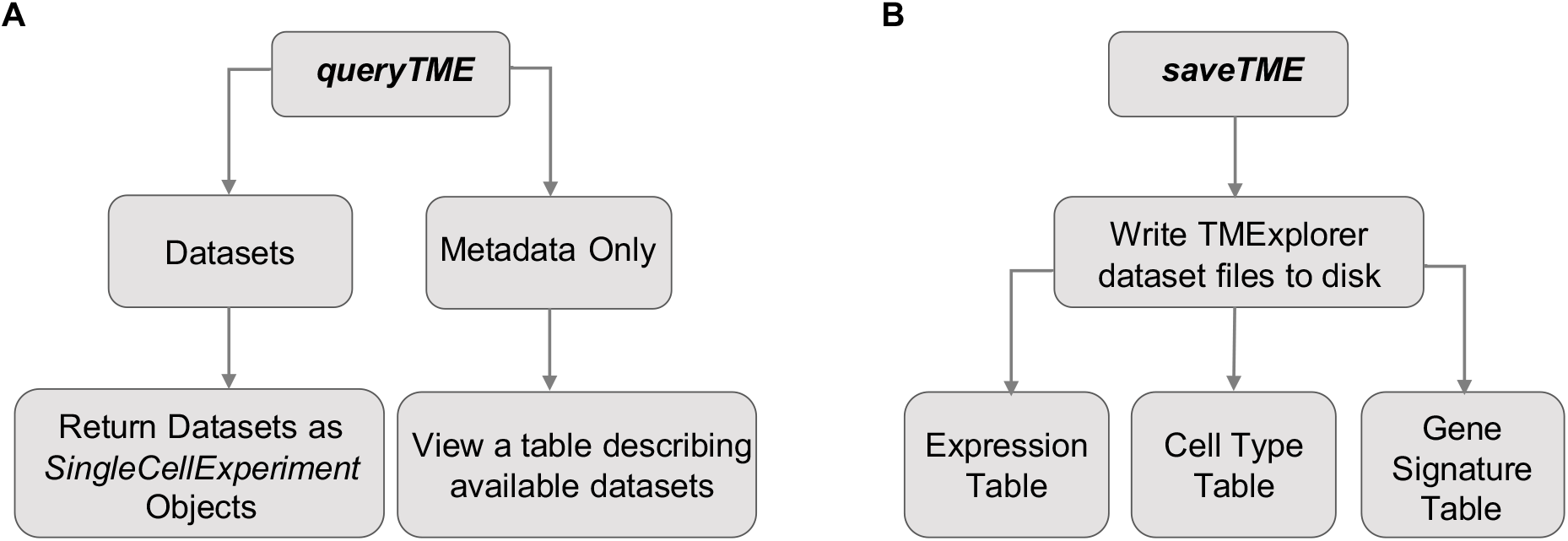
An overview of the main functions of TMExplorer. A) *queryTME* allows users to search and return datasets in either a descriptive table or as a list of *SingleCellExperiment* objects for analysis. B) *saveTME* allows users to write datasets to disk. For each dataset written to disk, up to three files are created; a table storing the expression data as either a CSV or matrix market file, depending on whether a dense or sparse matrix is passed to the function; a table containing the cells and their truth label, if available; and a table containing the cell type signature gene sets, if available.

After querying the database, a list of *SingleCellExperiment* objects are returned. The objects in this list can then be passed to any other algorithms that accept a *SingleCellExperiment* object, sparse *dgCMatrix*, or dense gene expression matrix for inclusion in a pipeline (Figure 3). Alternatively, the *saveTME* function can be used to write the returned data to disk for further manipulation or use in applications outside of R (Figure 2b). Figure 3 shows how *saveTME* can be used to save data for analysis in Python. In order to maintain consistency, the returned value is always a list of results, whether or not multiple datasets match the query.

**Figure 3:**
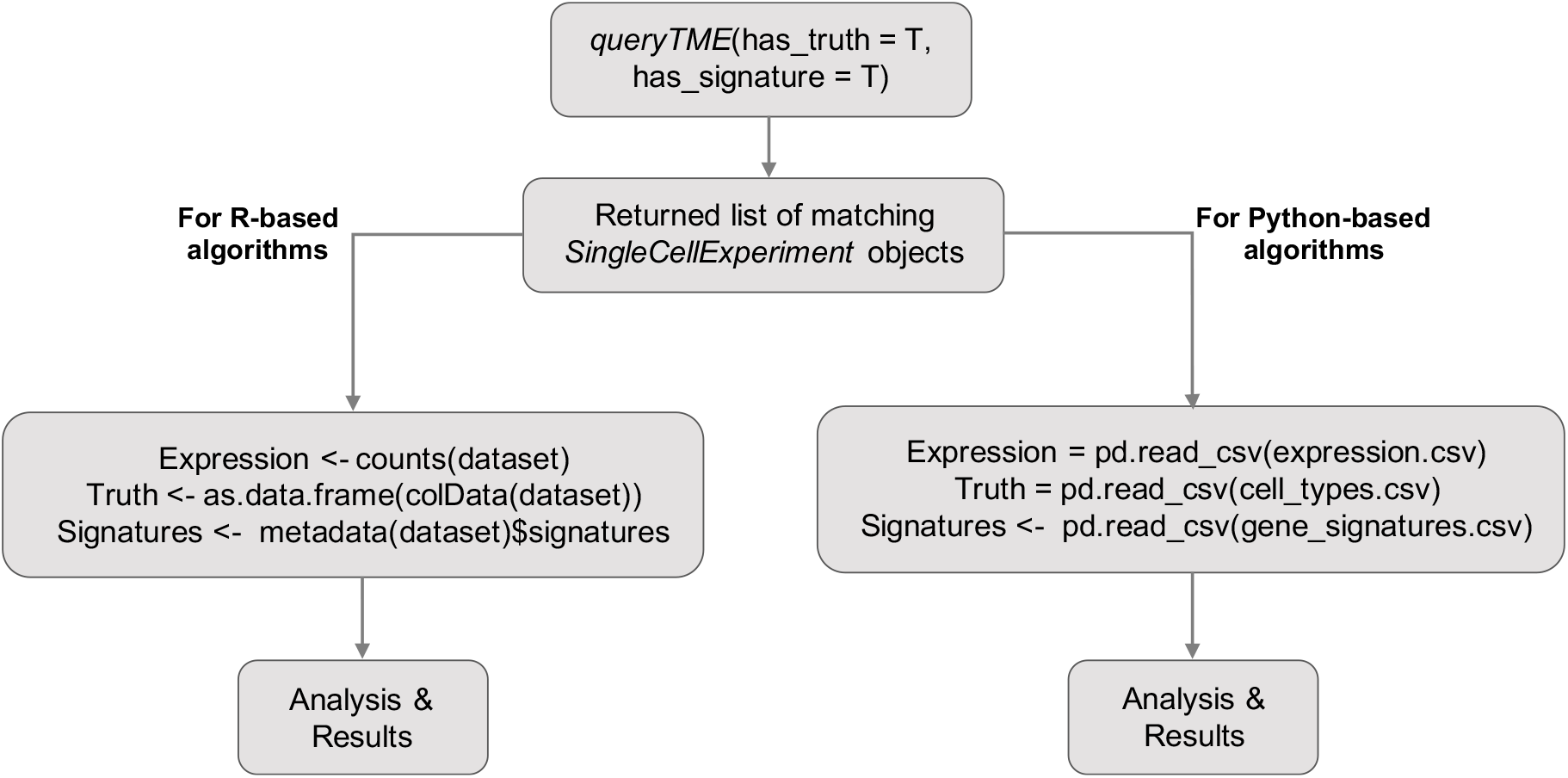
An example workflow of using TMExplorer to obtain datasets for the downstream analysis using Python and R. Users start by using *queryTME* to return all datasets that have cell type labels and cell type signature gene sets, which will get a list of matching datasets contained in *SingleCellExperiment* objects. Then, for R based algorithms, users can pass the *SingleCellExperiments* directly if that is supported, or users can pass the individual components required. For Python based algorithms, *saveTME* can be used to save the files for each dataset to disk, which can then be opened in Python for analysis.

### TMExplorer Database Contents

TMExplorer is a curated collection of TME scRNA-seq datasets that have been made available as an R-package. We created TMExplorer to improve accessibility and sharing of tumour scRNA-seq data. It acts as a single-entry point to various tumour scRNA-seq datasets for users interested in studying gene expression of the TME at the single-cell level. Currently, the collection contains 21 datasets, including 18 datasets derived from human tumours and 3 datasets derived from mouse tumours. This comprises 12 different cancer types including leukemia (22, 24), breast (2, 27, 28), colorectal (3), glioblastoma (7, 9, 31), glioma (23), head and neck (20), astrocytoma (21), oligodendroglioma (8), melanoma (1, 5, 19), lung carcinoma (6), pancreatic (25, 29) and prostate (26). Numbers of cells and genes vary across datasets and fall within the range of 5,796-57,915 genes and 74-57,530 cells. Each dataset is provided as processed gene expression data. We did not include raw scRNA-seq data (i.e. FASTQ files) in our collection because these files tend to be very large and can be accessed through the SRA, if available. Out of the 21 datasets, cell-type annotations are also provided for 12 datasets and gene signature information is provided for 9 datasets, so that users may access and use this information in their analyses. Users can browse through the available datasets using the metadata table and then choose which dataset(s) they would like to analyze. Users can also save the datasets for use outside of R, for instance in Python or web-based analysis pipelines.

### TMExplorer Search Capability

An important feature of TMExplorer is that it acts as both a database and search tool that can be easily implemented in one’s own workflow. Some other currently available scRNA-seq databases have a search function, but cannot be easily integrated into workflows because they are web-based (12–15). Currently available R-based scRNA-seq databases lack built in search tools, requiring users to access vignettes to see the available data before it can be retrieved for use in a pipeline (16). TMExplorer provides a search tool that allows users to search for datasets that fit their needs by tumour type, sequencing technology, source organism, and more (Table 1), all from the R command line. This makes TMExplorer an improvement over both R-based and web-based databases because users are able to browse and query data from the same console they are using for analysis. By including a search tool and database in a single package, TMExplorer provides a single point of entry to include TME scRNA-seq data into data analysis pipelines.

### Case Studies

In this section, we bring two example applications where TMExplorer can be used to facilitate data analysis. In the *case study 1*, we show how TMExplorer can be combined with automated cell-type identification algorithms to identify different cell types in TME scRNA-seq data. Here, we also show how users can return datasets with both the signature gene sets and gold standard annotations needed for testing cell-type identification. In *case study 2*, we show how TMExplorer can be integrated with the algorithms for inferencing copynumber variations in individual cells and facilitate the separation of malignant and non-malignant cells in multiple tumour scRNA-seq datasets of the same cancer type.

#### Case Study 1: Identifying different cell types in TME scRNA-seq data

Often, when using TME scRNA-seq data, we are interested in the cellular composition of the dataset. In order to find this, automated cell type identification algorithms are used. This is usually done by first clustering the cells, and then assigning appropriate cell type labels to each cluster (42). In Figure 4, we show how TMExplorer can be combined with a clustering method (e.g. Seurat (43)) and a cluster labelling method (e.g. GSVA (38)) to create a workflow for the identification of cell populations within a dataset. Seurat requires only the gene expression matrix to perform clustering, but GSVA requires a list of cell-type signature gene sets in addition to the expression matrix. TMExplorer can return all of the datasets that have signature gene sets available using *queryTME(has_signatures = TRUE)*. If after identifying the cell types within a dataset, users want to assess the performance of their workflow by comparing the automated annotations to those reported alongside the dataset, the *has_truth = TRUE* parameter can be added to *queryTME* to only return datasets that have gold standard labels available. Seurat and GSVA can be replaced by any other tool that accepts a *SingleCellExperiment* object or a matrix of gene expression values, providing flexibility for users to incorporate TMExplorer into their own workflows.

**Figure 4:**
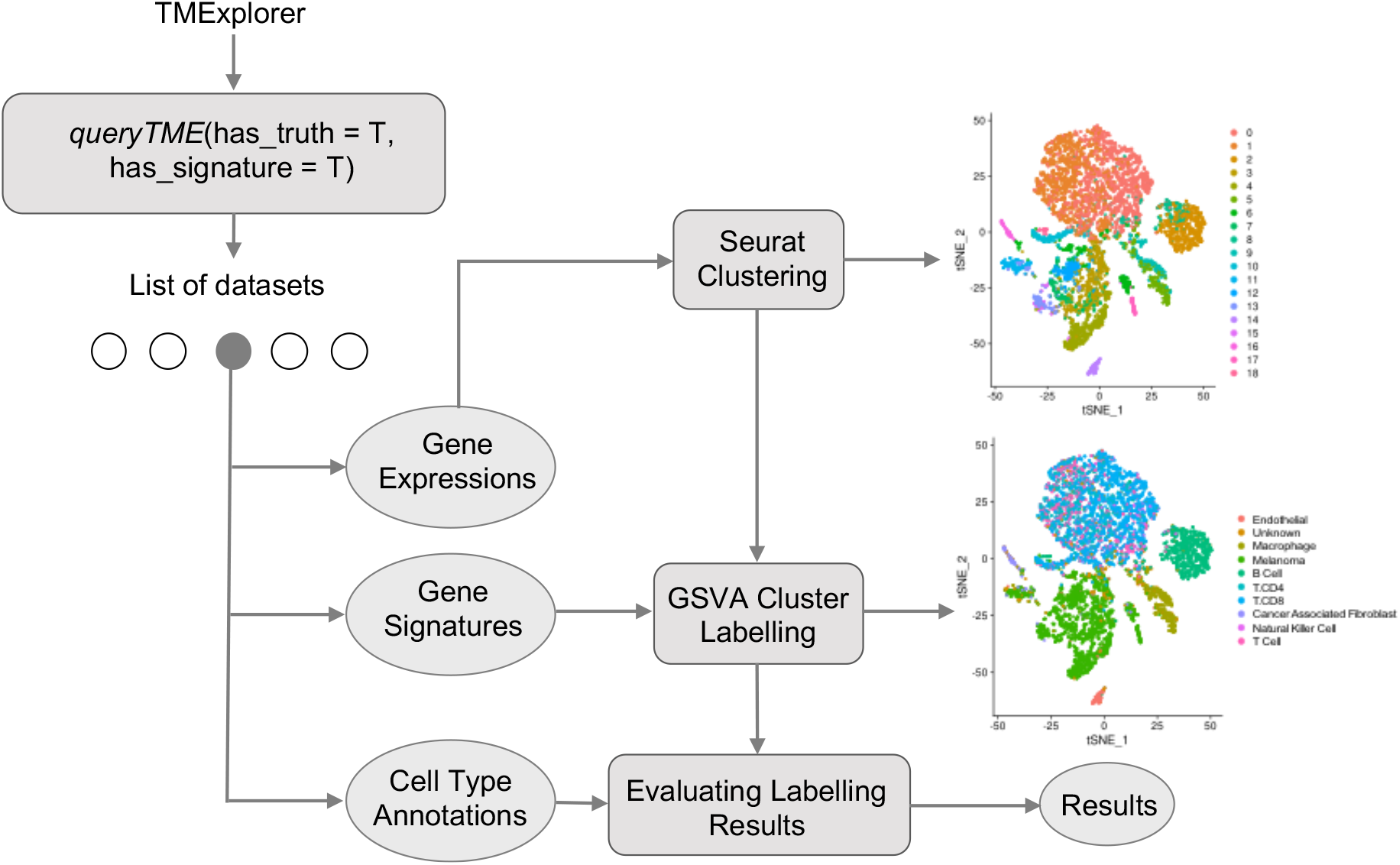
A case study on using TMExplorer to identify cell types. A case study showing how TMExplorer can be used to obtain datasets that can be used for cell cluster labelling via Seurat and GSVA. *queryTME* can be used to return those datasets which have both gene signatures and cell type annotations required for testing the automated identification of cell types. The expression data can be passed to Seurat for cell clustering, and the gene signatures can be used by GSVA to identify the cell types in Seurat’s clusters. Finally, the cell type annotations can be used as the truth labels to measure the performance of the results obtained by Seurat clustering followed by GSVA.

#### Case Study 2: Inferencing copy number variations in multiple datasets of the same cancer type

Single cell sequencing is an important tool that enables the dissection of TMEs into malignant and non-malignant cells. Researchers interested in comparing the tumour composition across different datasets of a specific cancer type would have to collect datasets from different sources prior to application of separation methods. With TMExplorer, users can easily access multiple datasets of a specific tumour type, as well as the accompanying cell type annotations and/or gene signature information from one location, thus avoiding inconsistencies when acquiring data from different databases. TMExplorer can be easily incorporated with other packages into workflows for the analysis of scRNA-seq data, therefore enabling users to access and use the data entirely within R.

Figure 5 displays an example workflow that uses *queryTME(tumour_type = “Glioblastoma”)* to retrieve datasets of a specific cancer type (i.e. glioblastoma) for use in the downstream analysis. In this example, we retrieved glioblastoma datasets from the TMExplorer database as *SingleCellExperiment* objects and converted them to gene expression count data matrices. We then applied a copy number variation (CNV) inferencing method called CONICSmat (44) to each of the datasets individually, and generated heatmaps displaying the inferred CNV patterns. This allowed us to separate malignant and non-malignant cells considering their long-range CNV patterns. The proportion of malignant and non-malignant cells and the patterns of CNV across the different datasets can then be compared.

**Figure 5:**
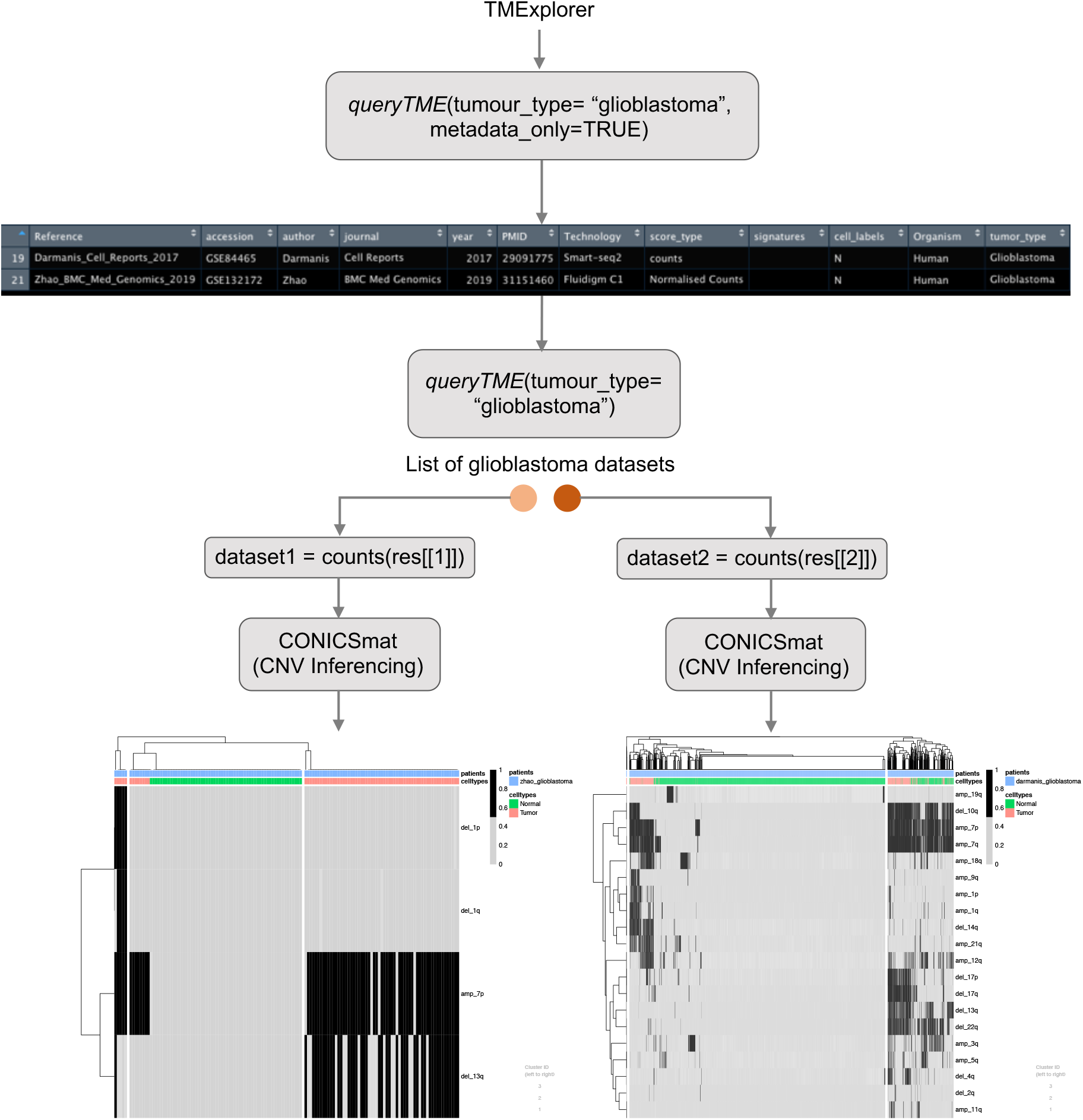
A case study on using TMExplorer for inferring CNVs. A case study showing how TMExplorer can be used to obtain multiple datasets for a specific tumour type, to be used with CNV-based separation methods, such as CONICSmat. *QueryTME* returns datasets of a specific tumour type, such as Glioblastoma. These datasets can then be inputted directly into large-scale CNV inferencing methods, such as CONICSmat.

## DISCUSSION

The emergence of single-cell RNA sequencing has enabled the study of tumour composition and phenotype. With the increasing use of scRNA-seq in cancer research, scRNA-seq data from TMEs continues to be generated and published. In order to streamline the data collection process for researchers interested in studying the TME, we created a curated database of TME scRNA-seq datasets, made available as an R-package called TMExplorer. Here we have built a database using a variety of cancers from multiple sources. We searched NCBI (11, 33) for TME scRNA-seq datasets that contain gene expression data, as well as comprehensive metadata such as tumour type, sequencing technology, cell type annotations, and gene signatures. In total, 21 datasets representing 12 different cancers are represented, along with their cell type annotations and cell type signature gene sets if they were available.

TMExplorer addresses a gap in currently available scRNA-seq databases by providing a focused, easily accessible database as an R package. TMExplorer has several advantages over other currently available scRNA-seq databases, the most prominent being:

1. Existing curated scRNA-seq databases consist of mostly normal tissue or non-cancer data and relatively few cancer datasets. To allow researchers to easily locate and access TME scRNA-seq data, we curated publicly available TME datasets and made them available in a database accessible as an R package. With TMExplorer, researchers can access all publicly available TME scRNA-seq datasets from a single location and can also return multiple datasets that match their desired criteria with a single command.
2. TMExplorer provides a variety of search parameters (Table 1) that can be used to return a subset of the available data that matches specific criteria. The parameters were designed so that users can search for matching datasets without having to first view a list of all available datasets, making it easier and faster to access data of interest.
3. The majority of existing scRNA-seq databases can only be accessed online as webbased tools and are not easily incorporated into pipelines for analysis of scRNA-seq data. Since many researchers use R or Python for their analyses, we chose to provide TMExplorer as an R-package so that it may be easily integrated into existing pipelines.
4. Some analyses require more than just gene expression information, and TMExplorer provides cell type annotations and cell type signature gene sets alongside gene expression matrices, where they are available. This facilitates the use of a wider range of analysis methods without requiring additional work from the researchers.

We plan to keep the package updated with new datasets as they are published. Users of the package doing their own novel research will have access to an issue template on Github where they can submit their data for inclusion. The interested users will need to provide their scRNA-seq data as raw counts or normalized data and the corresponding metadata. Since the package is open source, those same users can create a fork of the repository and build it from source with their own data for pre-publication work.

In summary, TMExplorer allows researchers to easily access, share and integrate TME scRNA-seq data into their own analysis pipelines. TMExplorer can be used to access data needed for the validation of new algorithms and to allow researchers interested in the tumour microenvironment to study specific types of cancer.

## SOFTWARE AND DATA AVAILABILITY

All included datasets are from free, public sources (11, 33) and all source code is freely available on Github (https://github.com/shooshtarilab/TMExplorer). Additionally, it is currently undergoing review for acceptance to BioConductor. Users wishing to have their data added to TMExplorer can open an issue at https://github.com/shooshtarilab/TMExplorer/issues with a link to their study and data along with a brief description and we will review it for inclusion.

## ACKNOWLEDGEMENTS

This work was supported by the research grants from Lawson Health Research Institute, the University of Western Ontario, and Ontario Institute for Cancer Research.

